# Molecular mechanism of tRNA binding by the *Escherichia coli* N7 guanosine methyltransferase TrmB

**DOI:** 10.1101/2022.10.29.514367

**Authors:** Sarah K. Schultz, Kieran Meadows, Ute Kothe

## Abstract

Among the large and diverse collection of tRNA modifications, 7-methylguanosine is frequently found in the tRNA variable loop at position 46. This modification is introduced by the TrmB enzyme, which is conserved in bacteria and eukaryotes. Complementing the report of various phenotypes for different organisms lacking TrmB homologs, we report here hydrogen peroxide sensitivity for the *Escherichia coli ΔtrmB* knockout strain. To gain insight into the molecular mechanism of tRNA binding by *E. coli* TrmB in real-time, we developed a new assay based on introducing a 4-thiouridine modification at position 8 of *in vitro* transcribed tRNA^Phe^ enabling us to fluorescently labelled this unmodified tRNA. Using rapid kinetic stopped-flow measurements with this fluorescent tRNA, we examined the interaction of wild-type and single substitution variants of TrmB with tRNA. Our results reveal the role of SAM for rapid and stable tRNA binding, the rate-limiting nature of m^7^G46 catalysis for tRNA release, and the importance of residues R26, T127 and R155 across the entire surface of TrmB for tRNA binding.

## Introduction

Transfer RNAs (tRNAs) contain the greatest density and diversity of modifications among RNA species (1). Among the assortment of chemical modifications, single methylations occur at different atoms of each of the four canonical nucleobases or the ribose sugar and are introduced by a collection of unrelated methyltransferases (2,3). Whereas some modifications are present in only certain tRNA isoacceptors in specific organisms, other modifications are highly conserved. The N7-methylguanosine (m^7^G) modification is common in bacterial and eukaryotic tRNAs, and has been found in a few archaea (4). Most often, m^7^G is present at position 46 in the variable loop of tRNAs, although it has additionally been found at tRNA positions 34, 36, and 49 in a handful of organisms (4). Within tRNAs, the base at position 46 is involved in a tertiary base pair with C13-G22, where it acts as a staple connecting the variable loop and the D arm. At this position vital for tRNA tertiary structure, the m^7^G46 modification introduces a site-specific positive charge amongst the negatively charged tRNA backbone (5,6). In *Escherichia coli*, m^7^G46 is introduced in about half of all tRNAs by TrmB, an S-adenosylmethionine (SAM)-dependent methyltransferase (Figure 1) (7,8).

**Figure 1.**
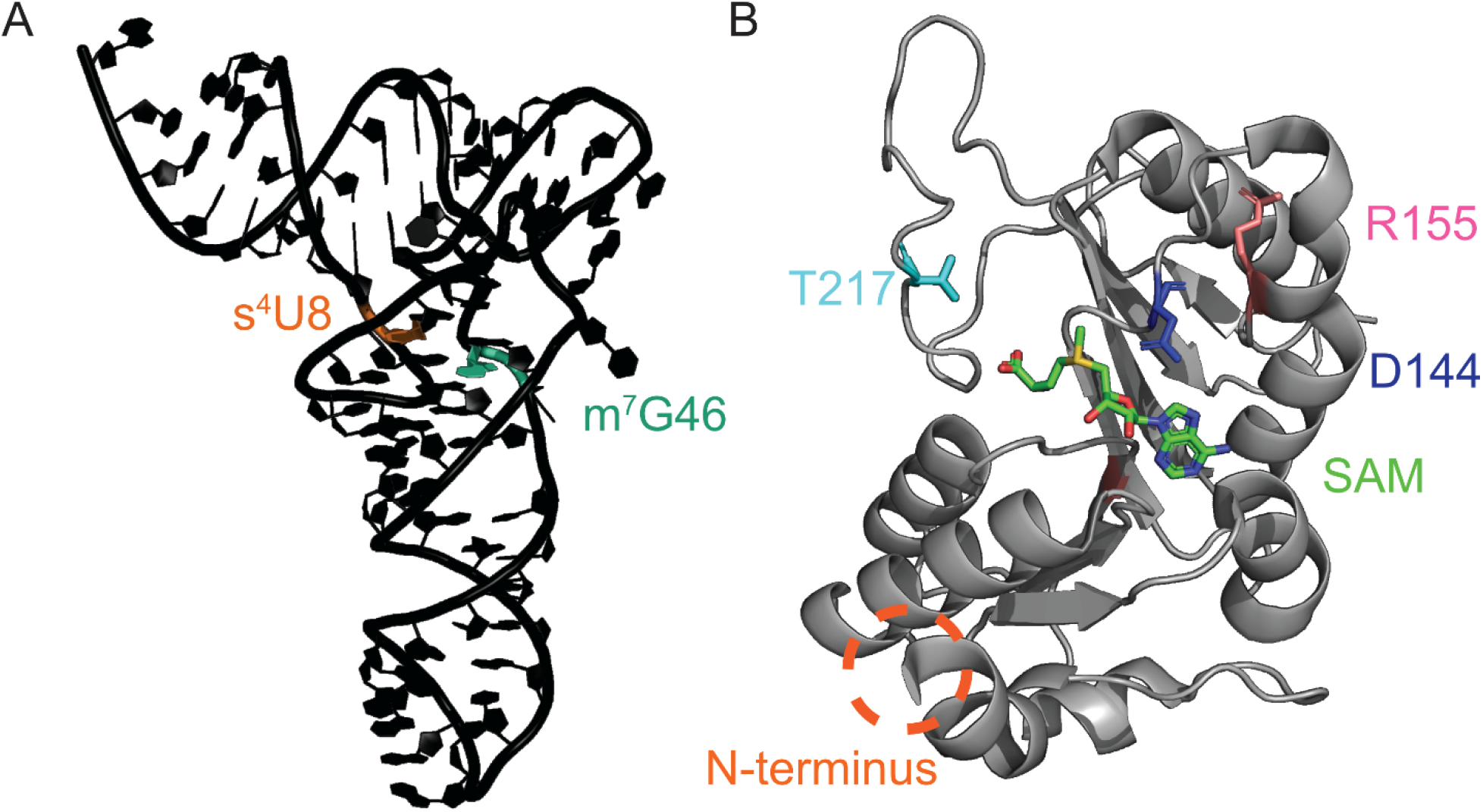
Structures of tRNA and TrmB. **(A)** Crystal structure of unmodified *E. coli* tRNA^Phe^ (PDB 3L0U; (51)) highlighting the locations of G46, where TrmB introduces the 7-methylguanosine modification (m^7^G, teal) and U8, which was modified by ThiI and IscS to 4-thioururidine (s^4^U, orange) for subsequent fluorescent labeling in this study. **(B)** Crystal structure of *E. coli* TrmB in complex with its cofactor SAM (PDB 3DXY; (8)). The methyldonor SAM is shown in green, and residues addressed in this study are shown as sticks, with residue D144 highlighted in blue, T217 in teal, and R155 in pink. As the first 36 amino acids of TrmB were not resolved in this structure, the location of the N-terminus, which would include residue R26, is circled in orange.

Similar to many tRNA modification enzymes, TrmB is non-essential in bacteria and knockout of the *E. coli trmB* gene does not impair growth under ideal conditions (7). While this observation has raised questions about the functions of TrmB and other tRNA modification enzymes, newer findings in different organisms suggest a role for the m^7^G46 modification under certain disease or stress conditions. Several recent studies have identified an importance for the human TrmB homolog METTL1 for cancer cell progression across several cancer types (reviewed in (9,10)), wherein the m^7^G46 modification prevents degradation of specific tRNAs, leading to an increase in global translation in addition to biased translation of growth-promoting genes (11-13). Thus, this tRNA methyltransferase has gained attention as a potential biomarker for cancer prognosis and as a potential chemotherapeutic target.

Moreover, human mutations resulting in the reduction of m^7^G methylation have been associated with a distinct form of microcephalic primordial dwarfism (14). Yeast lacking the TrmB homolog Trm8 display a specific growth defect in synthetic minimal media containing 2% glycerol at 38°C (15). This phenotype is exacerbated upon deletion of certain additional tRNA modification enzymes due to rapid decay of specific tRNA isoacceptors (16). Within eubacteria, the Gram-positive bacterium *Pseudomonas aeruginosa* lacking TrmB has been shown to be sensitive to hydrogen peroxide (H_2_O_2_) (17). A similar peroxide sensitivity has also been shown for the phytopathogenic fungus *Colletotrichum lagenarium* lacking the *trmB* homolog *aph1. C. lagenarium* cells lacking *aph1* additionally grow poorly in high salt concentrations and are unable to infect plant cells (18). Finally, *Thermus thermophilus ΔtrmB* cells exhibit severe growth defects at high temperatures, accompanied by decreases in other tRNA modifications, tRNA stability, and protein synthesis (19).

Some structural information about TrmB and its homologs is available, but we still do not know how exactly TrmB recognizes tRNA. Formation of m^7^G46 in bacteria requires only the TrmB protein; however, in eukaryotes, two non-related enzymes form a complex to modify tRNA. Whereas *E. coli* TrmB behaves as a monomer (7), other bacterial TrmB homologs have been shown to be homodimeric in solution, including *Bacillus subtilis, Streptococcus pneumoniae*, and *Aquifex aeolicus* TrmB homologs (8,20,21). In contrast, the eukaryotic TrmB homolog, known as Trm8 in yeast and METTL1 in humans, forms a heterodimer with Trm82 (yeast) or WDR4 (humans) (22,23). Crystal structures for *E. coli, A. aeolicus, B. subtilis*, and *Saccharomyces cerevisiae* TrmB homologs have been solved alone and/or with substrate SAM or product S-adenosylhomocysteine (SAH) (8,20,23-25). These structures have revealed that all TrmB homologs are class I methyltransferases deviating from the classic Rossman-fold only by an insertion between two β-strands (8,20). In *E. coli* TrmB, this insertion lacks secondary structure (8), whereas an α-helix is evident in other TrmB/Trm8 structures (20,23). A structure for any TrmB homolog in complex with tRNA does not yet exist and as such, we are lacking an understanding of how TrmB binds its substrate tRNA.

A prior study of *E. coli* TrmB identified several TrmB residues necessary for TrmB methylation activity *in vitro* but did not distinguish which of these residues are directly responsible for catalysis and which are involved in tRNA binding (26). Single-residue alanine substitutions at conserved residues R26, D144, H151, R154, R155, D180, T217, and E220 reduced methylation activity to less than ten percent of that of the wildtype enzyme, with substitutions at D144, R154, and R155 abolishing methyltransfer activity completely (26). Steady-state kinetic analysis for several equivalent single alanine substitutions in *A. aeolicus* TrmB has suggested that TrmB D144A, R155A, and T217A variants display slower methylation compared to wildtype TrmB, whereas SAM binding is unaffected for these variants (27). The authors of this study proposed a potential catalytic mechanism wherein D180 and the adjacent T179 residue (*E. coli* TrmB numbering) interact with the N1-proton and O6 atom of the target guanosine 46 base, respectively, thus positioning the N7 atom for nucleophilic attack by the SAM methyl group (27).

Here, we investigate the function of TrmB for *E. coli* fitness and dissect its interaction with tRNA using a novel assay. Thereby, we report that the model Gram-negative bacterium *E. coli* is sensitive to hydrogen peroxide stress in the absence of TrmB. In order to gain a better understanding as to how this tRNA modification enzyme binds tRNA, we prepared partially modified tRNA^Phe^ containing the 4-thiouridine 8 (s^4^U8) modification using purified ThiI and IscS enzymes. The reactivity of the s^4^U8 modification was then used to fluorescently label the tRNA for pre-steady-state kinetic analysis using a stopped-flow instrument. Moreover, to investigate the roles of specific TrmB residues for tRNA binding, we analyzed four inactive TrmB variants: R26A, D144A, R155A, and T217A. The binding of these variant enzymes to tRNA was examined in real-time using our stopped-flow assay, and the affinity of these enzymes for tRNA was determined using nitrocellulose filtration. Taken together, our results suggest that tRNA binding by TrmB is a complex, multi-step process aided by prior binding of SAM.

## Results

### E. coli ΔtrmB grows slowly in the presence of hydrogen peroxide

*E. coli trmB* is non-essential for cell growth and no growth phenotypes have been reported for the *E. coli ΔtrmB* strain under ideal growth conditions (7). Since previous work has shown the Gram-positive bacterium *P. aeruginosa* and the parasitic fungus *C. lagenarium* lacking the *trmB* gene grow slowly in media containing hydrogen peroxide, we asked whether *E. coli* lacking *trmB* displays the same phenotype (17,18). Indeed, *E. coli* BW25113 *ΔtrmB* reproducibly grows slower than its parental strain in LB medium supplemented with H_2_O_2_. Although the exponential and stationary phases are not affected by *trmB* knockout, the lag phase for the *ΔtrmB* cells is longer than that of the wildtype (Figure 2). Thus, an increased sensitivity to hydrogen peroxide in the absence of *trmB* seems to be common at least across Gram-positive and negative bacteria, in addition to the eukaryotic fungus *C. lagenarium*.

**Figure 2.**
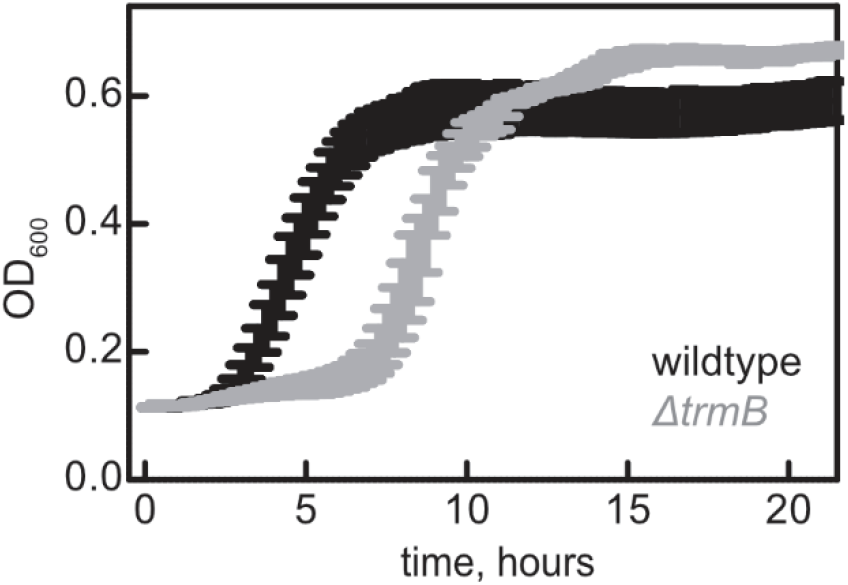
*E. coli* BW25113 *ΔtrmB* grows slowly in LB medium containing hydrogen peroxide. Four biological replicates for wildtype *E. coli* BW25113 (black) and *E. coli* BW25113 *ΔtrmB* (grey) were seeded with a starting OD_600_ of 0.1 in a 96-well plate. Average OD_600_ values recorded every fifteen minutes, and error bars represent standard error of the mean (SEM).

### Partial modification and fluorescent labeling of tRNA^Phe^ for rapid-kinetic analysis of tRNA binding by TrmB

In order to examine how TrmB is binding tRNA, we sought to introduce a fluorescent label into tRNA that would enable a fluorescence change that could be observed in real-time using stopped-flow fluorescence spectrometry. Within the three-dimensional tRNA structure, the s^4^U8 residue within the acceptor stem is in close proximity to the TrmB target nucleoside G46 (Figure 1A). Previous studies of tRNAs interacting with the ribosome have exploited the reactive nature of the s^4^U8 thiol modification to introduce different fluorescent dyes attached to iodoacetamide into native tRNA isoacceptors purified from cells (28). In order to study the interaction of TrmB with its substrate tRNA lacking m^7^G46 in addition to other modifications, we instead *in vitro* transcribed tRNA and introduced the single s^4^U8 modification using purified ThiI and IscS enzymes for fluorescent labeling with 5-iodoacetamidofluorescein (5-IAF) (Figure 3A).

**Figure 3.**
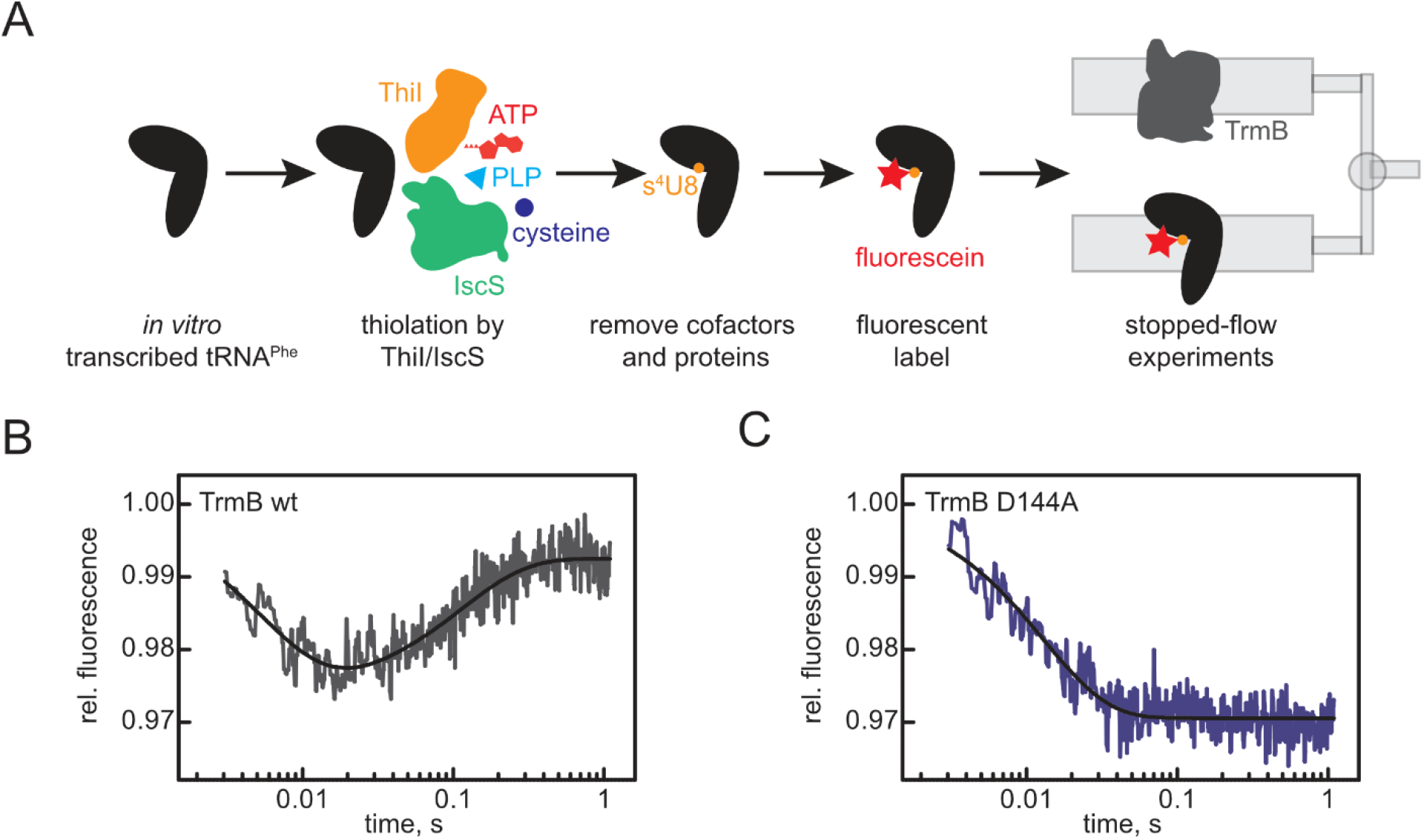
Partial modification and fluorescent labeling of tRNA^Phe^ for rapid-kinetic stopped-flow experiments. **(A)** Labeling scheme detailing the introduction of s^4^U8 by ThiI and IscS, modified tRNA purification, and fluorescent labeling with fluorescein for use in rapid kinetic assays. **(B)** Rapid mixing of 15 µM wildtype TrmB and 50 µM SAM with 1 µM fluorescein-s^4^U8-tRNA^Phe^.The data was fit to a 2-exponential equation to determine apparent rate constants: *k*_app1_: 180 s^−1^ and *k*_app2_: 9 s^−1^. **(C)** Rapid mixing of catalytically inactive 15 µM TrmB D144A and 50 µM SAM with 1 µM fluorescein-s^4^U8-tRNA^Phe^. Fitting the data with a 1-exponential equation determined a *k*_app1_ of 77 s^−1^.

Using the fluorescein-s^4^U8-tRNA^Phe^, we monitored the binding of wildtype TrmB to tRNA in the millisecond-to-seconds range using a stopped-flow apparatus. In the presence of 50 µM SAM, mixing 15 µM TrmB with 1 µM fluorescein-s^4^U8-tRNA^Phe^ resulted in a biphasic signal with an initial fluorescence decrease followed by a subsequent fluorescence increase (Figure 3B). This curve was fit with a 2-exponential equation, yielding apparent rates of ~180 s^−1^ (*k*_app1_) and ~9 s^1^ (*k*_app2_) (Figure 3B). In contrast, mixing fluorescein-s^4^U8-tRNA^Phe^ with the catalytically inactive variant TrmB D144A resulted in a single fluorescence decrease with an apparent rate of 77 s^−1^ (Figure 3C). As we will show below, this variant is deficient only in methylation, but not tRNA binding. In the presence of SAM, wildtype TrmB forms product (26,29). Thus, the fluorescence decrease reflects an event related to tRNA binding prior to methylation. Conversely, the fluorescence increase, that we observed only with the wildtype enzyme, is likely to occur at a step after catalysis and may reflect release of the methylated tRNA.

### TrmB variant binding to tRNA^Phe^

As previous work with *E. coli* TrmB variants has measured only tRNA methylation, but not tRNA binding, we investigated which TrmB residues are important for tRNA binding. To this end, we prepared two inactive TrmB variants that we hypothesized would retain tRNA binding ability while being unable to methylate tRNA: TrmB D144A and T217A (Figure 1B). Both residues are proposed to participate in methylation, wherein the side chain of the aspartate equivalent to D144 has been shown to form hydrogen bonds with N6 of the SAM adenosine moiety in *B. subtilis* TrmB (25). Residue T217 is present within the TrmB-specific insertion that is unstructured in *E. coli* TrmB (8). Within *B. subtilis* TrmB, this threonine residue is located at the bottom of the SAM-binding pocket, wherein its side chain hydrogen bonds to the SAM methionyl moiety (20,25). However, this interaction is not observed within the *E. coli* TrmB–SAM complex (8) and thus T217 may only move in close enough proximity to interact with SAM upon structural rearrangement after tRNA binding. Alternatively, this residue may play a different role in *E. coli* TrmB.

To measure the affinity of each TrmB variant for tRNA^Phe^, nitrocellulose filter binding using tritium-labeled tRNA was performed. Previously, we have shown that wildtype TrmB in the absence of SAM has a relatively low affinity for tRNA with a dissociation constant of 6.6 µM (Figure 4A, Table 1) (29). When TrmB wildtype is pre-incubated with SAM, the affinity of TrmB for tRNA increases about three-fold to 2.1 µM (Figure 4A, Table 1). Thus, prior binding of SAM to TrmB has a positive allosteric effect on subsequent tRNA binding, as has previously been shown for TrmA (30).

**Figure 4.**
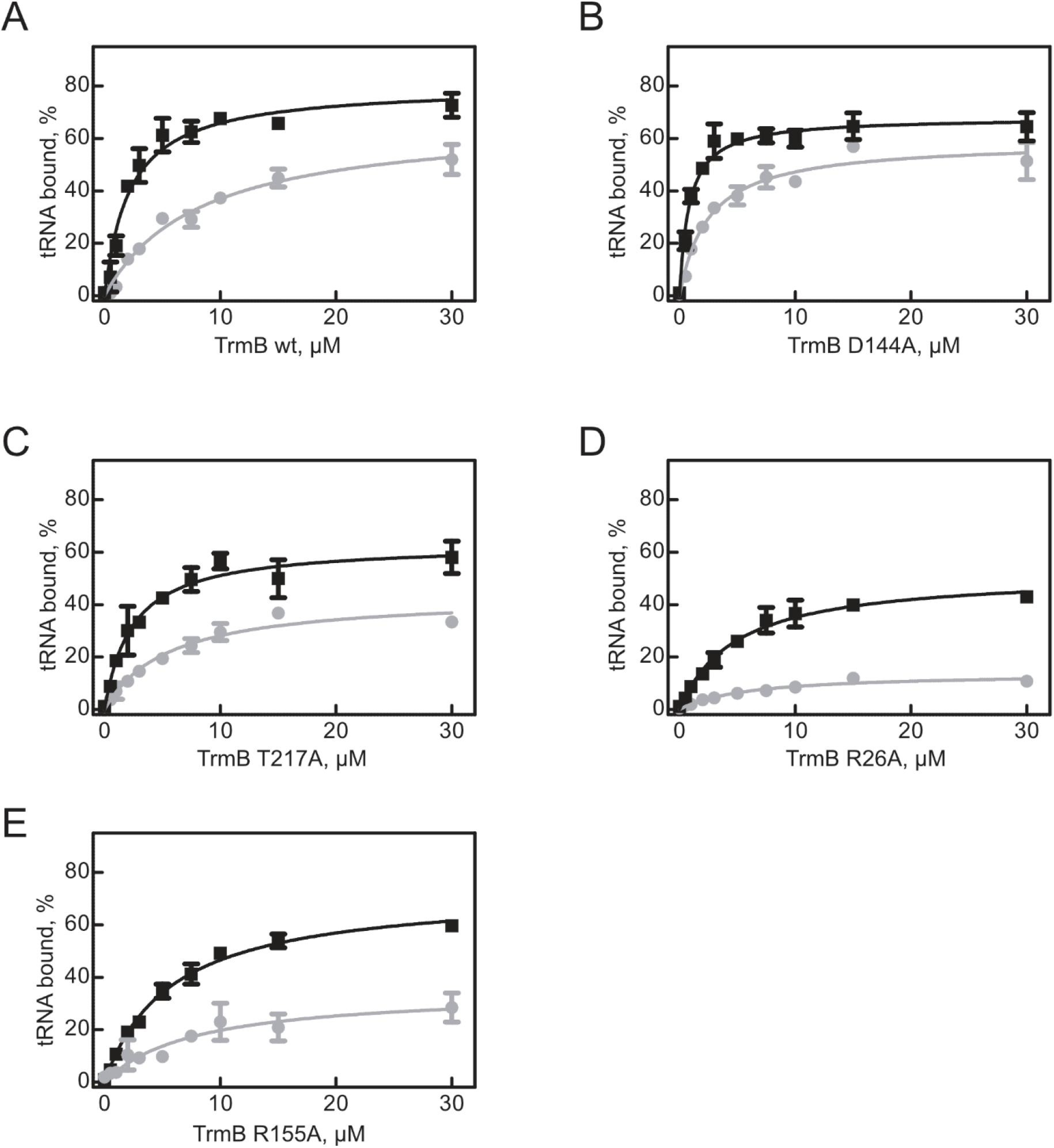
tRNA binding by TrmB wildtype and variants. To determine the affinity of TrmB in the presence (black squares) or absence (grey circles) of 50 µM SAM, 20 nM of [^3^H]tRNA^Phe^ was incubated with increasing concentrations of TrmB wildtype **(A)**, TrmB D144A **(B)**, TrmB T217A **(C)**, TrmB R26A **(D)**, or TrmB R155A **(E)**. Percent tRNA bound was determined by nitrocellulose filtration. Averages of at least three experiments are shown with error bars representing standard deviation. The data was fit with a hyperbolic equation to determine the dissociation constants (see Table 3).

**Table 1.**
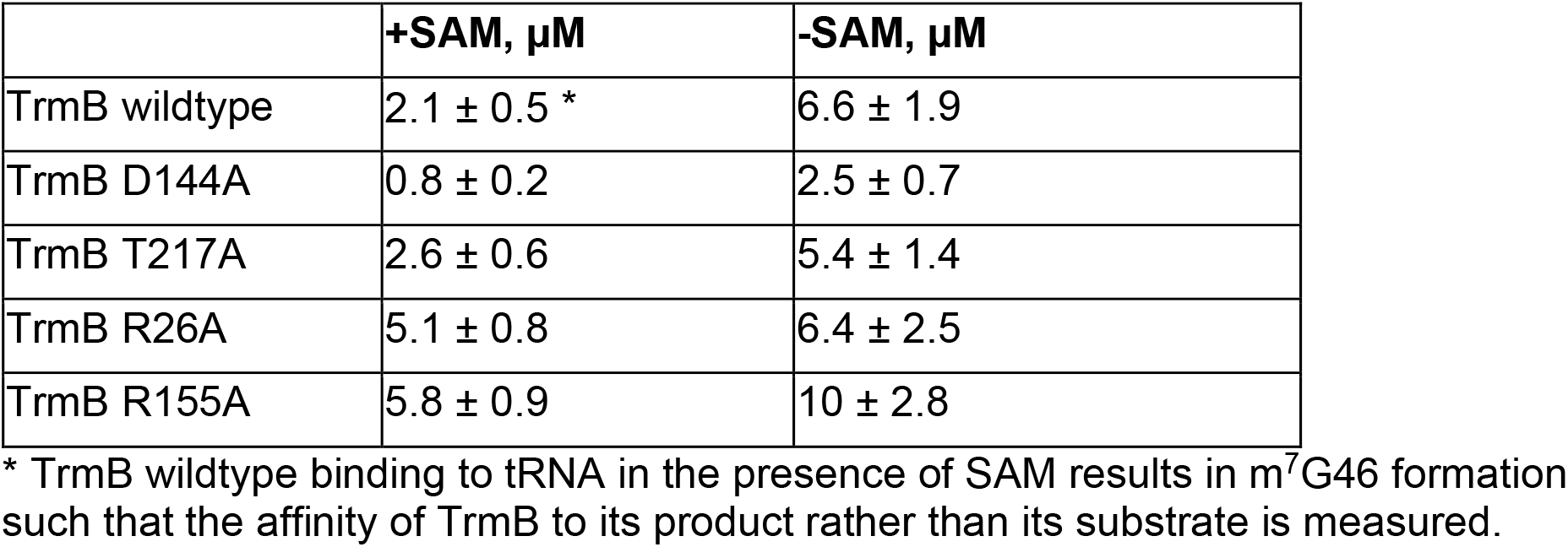
Average dissociation constants (K_D_) for TrmB wildtype and variants binding to [^3^H]tRNA^Phe^ in the presence and absence of 50 µM SAM.

Next, we examined the binding of inactive TrmB variants to tRNA^Phe^. In both the absence and presence of SAM, TrmB D144A binds tRNA with a 2.5-fold higher affinity than the wildtype enzyme, with a *K*_D_ of 2.5 µM and 0.8 µM, respectively (Figure 4B, Table 1). Unlike TrmB wildtype, this inactive TrmB variant cannot form product during the course of the reaction and thus its tRNA affinity is reflective of binding unmodified substrate, rather than m^7^G-containing product. This difference in binding unmodified tRNA rather than modified tRNA likely explains the higher affinity of TrmB D144A for tRNA compared TrmB wildtype when SAM is present. Since the other TrmB variants discussed below also do not form product (26), their affinities for tRNA will be compared to that of catalytically inactive TrmB D144A. Accordingly, we observed that alanine substitution at residue T217 reduces tRNA affinity two-to-three-fold in both the presence and absence of SAM compared to TrmB D144A (Figure 4C, Table 1).

Additionally, we prepared two inactive TrmB variants that we hypothesized would be deficient in tRNA binding: R26A and R155A (Figure 1B). Residue R26 is present within the N-terminal extension of TrmB that is presumably flexible because this region was not resolved in the *E. coli* TrmB structure (8). As a positively charged residue, R26 may contribute to tRNA binding via electrostatic interactions. Another positively charged residue, R155, is one of two adjacent conserved arginine residues along with R154. Previous work showed that alanine substitution of either residue abolishes methylation activity (26).

In the absence of SAM, TrmB R26A has a 2.5-fold lower affinity for tRNA compared to the catalytically inactive TrmB D144A variant, whereas in the presence of SAM, the affinity of TrmB R26A is even more affected, with a six-fold higher *K*_D_, at 5.1 µM (Figure 4D, Table 1). Notably, the maximum binding amplitude for TrmB R26A in the absence of SAM is very low, with only around 20% tRNA binding, further suggesting that this TrmB variant is deficient in stable tRNA binding (Figure 3D). In the absence of SAM, TrmB R155A binds tRNA with a *K*_D_ of 10 µM, which is about four-fold lower than the affinity of TrmB D144A for tRNA (Figure 3E, Table 1). Similarly, in the presence of SAM, the affinity of TrmB R155A is about seven-fold lower compared to the catalytically inactive variant (Figure 4E, Table 1). Like the R26A variant, the maximal level of tRNA bound by TrmB R155A is low, reaching only about 30% tRNA bound (Figure 4E). Thus, substitution at residues R26 and R155 negatively affect tRNA binding.

### Rapid kinetic analysis of inactive TrmB variants

To monitor the interaction between TrmB variants and tRNA in real-time, we repeated the stopped-flow assays using fluorescein-s^4^U-tRNA^Phe^ with TrmB variants. Binding of the TrmB D144A variant to tRNA in the absence of SAM (Figure 5A) displays a similar single-phase fluorescence decrease as seen when mixing this variant and fluorescein-s^4^U8-tRNA^Phe^ in the presence of SAM (Figure 3B). Fitting the binding curve of TrmB D144A in the absence of SAM with an exponential equation yielded an apparent rate of about 60 s^−1^ which is comparable to the apparent rate of 77 s^−1^ observed in the presence of SAM.

**Figure 5.**
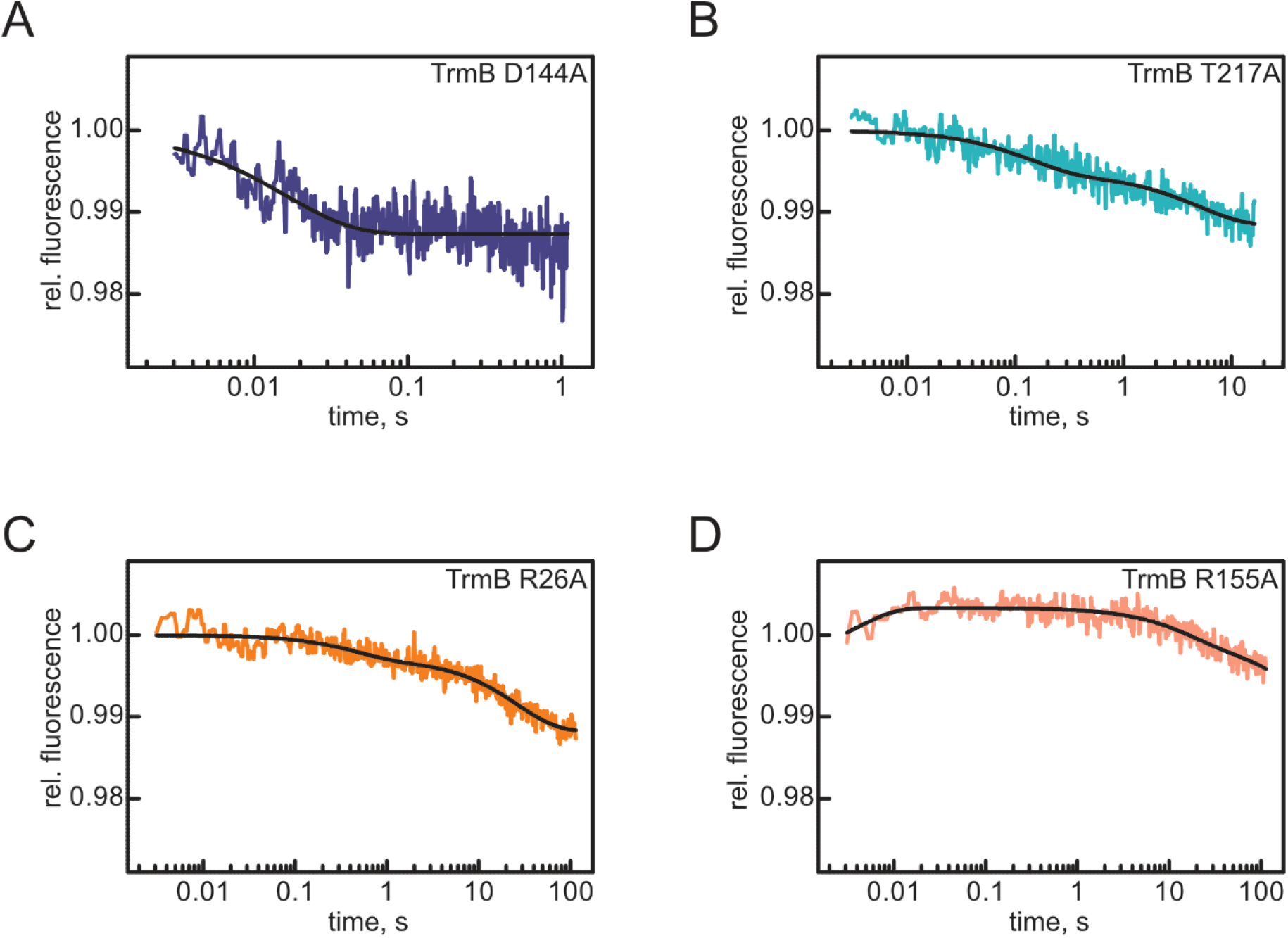
Rapid kinetic analysis of TrmB variant binding to tRNA. Time courses displaying rapid mixing of 15 µM TrmB variants with 1 µM fluorescein-s^4^U8-tRNA^Phe^ in a stopped-flow apparatus. Each trace is an average of at least eight independent time courses. Note the different x-axis range between panels. The time course for TrmB D144A **(A)** was fit to a 1-exponential equation with a *k*_app1_ of 62 ± 5 s^−1^, whereas data for TrmB T217A **(B)** was fit to a 2-exponential equation with a *k*_app1_ of 8 ± 1 s^−1^ and a *k*_app2_ of ~0.2 s^−1^. Time courses for TrmB R26A **(C)** were fit to a 2-exponential equation, with a *k*_app1_ of 2 ± 0.2 s^−1^ and *k*_app2_ of ~0.04 s^−1^. Finally, data for TrmB R155A **(D)** was fit to a 3-exponetial equation, revealing a *k*_app1_ of 176 ± 46 s^−1^, *k*_app2_ of 2 ± 1 s^−1^, and a *k*_app3_ of ~0.03 s^−1^.

The affinity of the inactive TrmB T217A variant is similar to that of the wildtype enzyme, but reduced compared to TrmB D144A (Figure 4, Table 1). Accordingly, the fluorescence change observed upon mixing TrmB T217A with fluorescein-s^4^U8-tRNA^Phe^ is different from TrmB wildtype and D144A. Here, we observe two subsequent fluorescence decreases, with apparent rates of 8 s^−1^ (*k*_app1_) and 0.2 s^−1^ (*k*_app2_) (Figure 5B).

Likewise, mixing TrmB R26A with fluoresceine-s^4^U8-tRNA^Phe^ reveals two fluorescence decreases with a *k*_app1_ of 2 s^−1^ and *k*_app2_ of 0.03 s^−1^ (Figure 5C). Finally, the binding of TrmB R155A with fluoresceine-s^4^U8-tRNA^Phe^ in real time occurs in three phases with an initial fluorescence increase (*k*_app1_) with a rate of 176 s^−1^, followed by two decreases in fluorescence with rates of 3 s^−1^ (*k*_app2_) and 0.03 s^−1^ (*k*_app3_) (Figure 5D). Thus, our rapid kinetic assays reveal fast tRNA binding in the case of the wildtype enzyme and the catalytically inactive variant TrmB D144A. In accordance with the defects in tRNA binding for TrmB T217A, R26A, and R155A compared to the other inactive TrmB variant D144A observed in equilibrium binding conditions in the nitrocellulose filtration assays (Figure 4, Table 1), the TrmB T217A, R26A, and R155A variants display vastly different tRNA binding time courses, suggesting that substitutions to these residues significantly perturb how these variants bind tRNA. (Figure 5).

### Rapid-kinetic analysis of catalytically inactive TrmB D144A binding to tRNA^Phe^

In order to kinetically characterize the association of TrmB to tRNA^Phe^ in the absence of methylation, we used the catalytically inactive TrmB D144A variant that is deficient in catalysis but not tRNA binding (Figure 4, 5, (26)). Increasing concentrations of TrmB D144A were rapidly mixed with fluorescein-s^4^U8-tRNA^Phe^ and one-exponential equations were fit to the resulting curves. In the absence of the methyldonor SAM, the apparent rate is not dependent on enzyme concentration, with an average rate of 69 ± 5 s^−1^ (Figure 6A). This suggests that in the absence of SAM the observed apparent rate is indicative of a unimolecular conformational change, rather than a bimolecular binding event. In contrast, when TrmB D144A is pre-bound to SAM, the apparent rate of tRNA binding is concentration-dependent, and therefore is indicative of a bimolecular interaction. Determination of the slope reveals an association rate constant, *k*_1_, of 2.4 ± 0.5 µM^−1^ s^−1^ (Figure 6B). As we will further discuss below, these results can be explained if a conformational change in TrmB is required for stable binding of tRNA.

**Figure 6.**
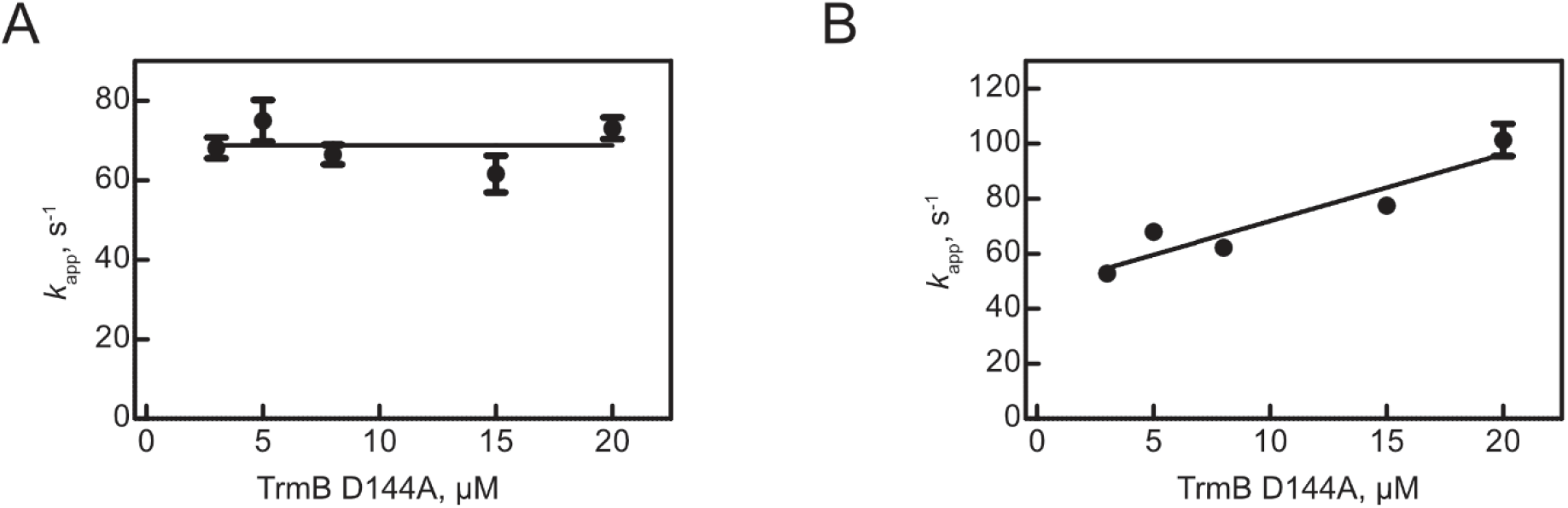
Concentration dependence of tRNA binding by catalytically inactive TrmB D144A. Time courses of TrmB D144A binding to 1 µM fluorescein-s^4^U8-tRNA^Phe^ were fit to single exponential equations to determine the concentration dependence of apparent rates. In the absence of SAM **(A)**, the *k*_app_ is not concentration-dependent, and the horizontal line represents the average apparent rate of 69 ± 5 s^−1^. In the presence of SAM **(B)**, the apparent rate of tRNA binding by TrmB is dependent on TrmB concentration. Thus, data was fit with a linear regression to determine the slope representing the association rate constant (*k*_1_) of 2.4 ± 0.5 µM^−1^ s^−1^.

## Discussion

### Peroxide sensitivity of E. coli ΔtrmB

In this study, we report for the first time a growth phenotype for an *E. coli ΔtrmB* strain, wherein *ΔtrmB* cells grow slowly in hydrogen peroxide-containing LB medium. This phenotype matches previous reports of phenotypes for *trmB* knockouts in *P. aeruginosa* and *C. lagenarium* (17,18). During the course of our study, this phenotype was reported to be absent for the *E. coli* BW25113 *ΔtrmB* strain, even in the presence of 10 mM H_2_O_2_ (31). However, this phenotype is highly reproducible in our experiments (Figure 2), and even the BW25113 wildtype strain does not significantly grow in the presence of 10 mM H_2_O_2_ within the small volume of 96-well plates (data not shown). This lack of growth for the BW25113 strain at high H_2_O_2_ concentrations has recently been corroborated by others (32). Given the unstable nature of hydrogen peroxide rendering it difficult to use precise concentrations and the conserved nature of this phenotype across different organisms, we conclude that *trmB* is important for fitness during oxidative stress.

It is notable that sensitivity to hydrogen peroxide stress is present in both Gram-positive and Gram-negative bacteria, in addition to *C. lagenarium*, a eukaryote. This suggests the m^7^G46 modification and/or TrmB plays a conserved role in mediating oxidative stress. Within *P. aeruginosa*, TrmB was found to mediate the peroxide stress response by optimizing translation of Phe and Asp codons, which are enriched within the *P. aeruginosa* catalase genes (17). The precise mechanism behind why TrmB enhances Phe/Asp translation remains unclear. It could be that the m^7^G46 modification is directly important for tRNA aminoacylation or for the role of tRNA on the ribosome during translation, or m^7^G46 could act to increase the cellular stability of certain tRNAs, thereby increasing their cellular abundances and availability for translation. Considering the demonstrated importance of the m^7^G46 modification by Trm8/Trm82 for preventing rapid decay of tRNA^Val^_(AAC)_ in yeast (16), in addition to the recent evidence that m^7^G46 formation by METTL1/WDR4 in mammals is important for the stability of at least tRNA^Arg^_(UCU)_ (11) or tRNA^Lys^_(CUU)_ (33) and tRNA^Lys^_(UUU)_ (13), it is likely that methylation by TrmB is important for the cellular stability of certain tRNAs in bacterial cells. This possibility will be interesting to examine in the future. Moreover, given the specific importance of m^7^G46 for only certain tRNAs across different organisms, investigation of the mechanism underlying why TrmB is important for peroxide tolerance in *E. coli* and whether or not this is similar to the determined mechanism for *P. aeruginosa* is interesting to consider.

### In vitro partial modification and fluorescent tRNA labeling

Fluorescent labeling of tRNAs at different specific internal sites by taking advantage of the reactivity of native tRNA modifications has been widely used to study the dynamics of tRNAs on the ribosome during protein translation (28,34). However, few studies have prepared *in vitro* transcribed tRNAs with only a select modification for internal fluorescent modification of a mostly unmodified tRNA (35,36). Using *in vitro* transcribed tRNA is advantageous for studying its interaction of tRNA modifying enzymes because this allows for examination of the binding of a modification enzyme to its substrate (rather than product), unmodified tRNA, without the laborious process of extracting specific tRNA isoacceptors from an appropriate knockout strain. Moreover, previous work has suggested TrmB homologs tend to modify tRNA in the relatively early stages of tRNA maturation (37,38), and *E. coli* TrmB seems to not have a preference for tRNA modification status (29). This suggests that a mostly unmodified tRNA is an appropriate substrate for studying the interaction between TrmB and tRNA. As ThiI has few sequence constraints for modifying U8-containing tRNAs and instead recognizes the overall L-shaped structure (39), this strategy can be used to introduce s^4^U8 within a wide selection of tRNAs. Thus, the here described preparation of fluorescently labelled, partially-modified tRNA for kinetics studies lays the foundation for study of additional tRNA modification enzymes or other tRNA-interacting enzymes, demonstrating another use for *in vitro* modified tRNAs (40).

### Molecular mechanism of E. coli TrmB interacting with tRNA

As we have previously shown, in comparison to other tRNA modification enzymes, *E. coli* TrmB has a relatively low affinity for tRNA with a *K*_D_ of about 6 µM in the absence of its cofactor SAM (Figure 4, Table 1) (29). Interestingly, *E. coli* TrmB binds tRNA with a lower affinity compared to its characterized homolog. Homodimeric *B. subtilis* TrmB was shown to bind tRNA^Phe^ in the absence of SAM with an affinity of about 0.1 µM using fluorescence anisotropy (25). This raises the possibility that TrmB enzymes with different quaternary structures may have distinct tRNA binding mechanisms. As such, the affinity may increase for homodimeric enzymes compared to monomeric *E. coli* TrmB. No investigations concerning the affinity of the eukaryotic Trm8/Trm82 heterodimer for tRNA have been conducted to date, and it is possible the heterodimeric complex again binds tRNA differently compared to monomeric bacterial TrmB. Supporting this possibility, differences in tRNA structural requirements have been identified between homodimeric *A. aeolicus* TrmB and heterodimeric yeast Trm8, wherein both enzymes require the T stem for modification activity, but only yeast Trm8 additionally requires the D stem (41,42). Moreover, the R26 residue, that we demonstrate here to be important for tRNA binding (Table 1, Figure 5), is not conserved in *B. subtilis* or *S. cerevisiae* TrmB homologs, and the N-terminal region present in mesophilic TrmB proteins is entirely absent in *A. aeolicus* TrmB. Notably, TrmB is one of several tRNA modification enzymes that utilizes only a single subunit in bacteria but requires an auxiliary protein for catalysis in eukaryotes (43,44). As such, a similar difference in tRNA affinity may exist for other bacterial tRNA modification enzymes and their eukaryotic two-subunit homologs.

Interestingly, we are demonstrating here that the affinity of TrmB for tRNA is significantly increased in the presence of SAM. This observation holds true for both wildtype TrmB, which forms m^7^G46 thus reporting product tRNA binding, as well as for the catalytically inactive variant TrmB D144A, which binds unmodified substrate tRNA (Table 1). Thus, prior binding of the cofactor SAM has a positive allosteric effect on the interaction of TrmB with tRNA, presumably by ordering and pre-orienting the active site. Previously, we have described a similar positive allostery for SAM and tRNA binding for the methytransferase TrmA responsible for the m^5^U54 modification in tRNAs (30).

In order to characterize the molecular mechanism of TrmB’s interactions with tRNA, we have established here a novel fluorescent assay enabling rapid-kinetic studies of tRNA binding to TrmB (Figure 3A). In the stopped-flow assay, we observe how fast TrmB binds to unmodified substrate tRNA giving us insight into the initial steps of TrmB’s molecular mechanism. Thus, the stopped-flow assay provides different type of information (kinetics of substrate binding) about TrmB wildtype than the nitrocellulose filtration assay (affinity of product binding). By comparing how tRNA interacts with wildtype TrmB compared to catalytically inactive TrmB D144A, we can gain insight into the kinetic mechanism of tRNA binding for this tRNA modifying enzyme. In these rapid-kinetic stopped-flow experiments, we observe first a fluorescence decrease when either wildtype TrmB or TrmB D144A interact with fluorescein-s^4^U8-tRNA in the presence of SAM. This fluorescence decrease upon tRNA binding may reflect partial unfolding of the tRNA structure as TrmB accesses its target base, G46, which is buried within the tRNA elbow forming a base pair triple with C13-G22. Notably, the apparent rate for the fluorescence decrease upon tRNA binding in the presence of SAM is two-fold higher for wildtype TrmB compared to TrmB D144A (Figure 3), suggesting that substitution of D144 for alanine slightly slows the association of TrmB with unmodified tRNA relative to wildtype TrmB.

Following the first fluorescence decrease in the rapid-kinetic stopped-flow assays with wildtype TrmB in the presence of SAM discussed above, we then observe a second fluorescence increase with an apparent rate of ~9 s^−1^. Presumably, this phase reflects the release of the modified tRNA, as this fluorescence change is absent when the catalytically inactive variant is mixed with tRNA in the presence of SAM (Figure 3C). Supporting this interpretation, the apparent rate for this step (measured at 20°C) is within the same order of magnitude as the *k*_cat_ for *E. coli* TrmB which has been reported to be 3.6 s^−1^ at 37°C (26). Thus, catalysis of m^7^G46 formation may be the rate-limiting step in the molecular mechanism of TrmB which is followed by rapid tRNA release. Notably, we have likewise reported that many tRNA pseudouridine synthases are characterized by a rate-limiting catalytic step (45).

To gain further insight into the mechanism of tRNA binding by TrmB, we performed a titration of TrmB D144A binding to tRNA in the stopped-flow experiments (Figure 6). In the absence of SAM, we observed that the apparent rate, *k*_app_, is not dependent on enzyme concentration, suggesting a unimolecular conformational change rather than a bimolecular binding event. In contrast, *k*_app_ was found to be dependent on TrmB D144A concentration in the presence of SAM, which suggests that in this case we observe a bimolecular binding event. This different kinetic behavior in the presence and absence of SAM indicates that different mechanisms are at play. As prior SAM binding has a positive allosteric effect on tRNA binding (vide supra, Table 1), it is conceivable that SAM binding may be necessary to induce a conformational change in TrmB, which structurally prepares TrmB for stable tRNA binding. As this conformational change has already occurred when TrmB is preincubated in the presence of SAM, the initial fluorescence decrease in the stopped-flow experiments likely reflects the bimolecular interaction of TrmB and tRNA. However, in the absence of SAM, it is likely that tRNA binding is rate-limited by a preceding unimolecular conformational change in TrmB which is slow due to the absence of SAM. Thus, the observed apparent rate is not dependent on the TrmB concentration. Thereby, our rapid-kinetic stopped-flow experiments provide further insight into the role of SAM for facilitating tRNA binding to TrmB.

### Structural features of tRNA binding by TrmB

In order to identify *E. coli* TrmB residues with roles in tRNA binding, we prepared four TrmB variants and for characterization both by equilibrium filter binding assays as well as our novel rapid-kinetic stopped flow assay. For the first time, this allows us to quantitatively investigate tRNA binding by TrmB independent of its catalytic activity allowing us to dissect the contribution of specific TrmB resides for these steps. As discussed, the catalytically inactive TrmB D144A variant binds tRNA tighter than TrmB wildtype because it reflects binding of unmodified substrate rather than modified product tRNA. Nevertheless, the rapid-kinetic stopped-flow experiments indicate that the D144A substitution may also slightly impair the kinetics of tRNA association (Figure 3). In comparison to TrmB D144A, the TrmB T217A variant has an approximately three-fold lower affinity for tRNA as measured in equilibrium nitrocellulose filtration binding (Table 1). Interestingly, the rapid-kinetic stopped-flow experiments provide further detailed insight into the mechanism of tRNA binding by TrmB variants. Here, we observed that tRNA associates to TrmB T217A much slower than to TrmB D144A (Figure 5).

Together, the equilibrium and rapid-kinetic experiments thus demonstrate unambiguously that TrmB T217A is not only impaired in m^7^G46 formation as previously reported, but that the substitution of T217 with alanine also impairs tRNA binding to TrmB. Similarly, even stronger defects with respect to the affinity and kinetics of tRNA binding were observed for TrmB R26A and TrmB R155A (Table 1, Figure 5). For both variants, the tRNA affinity in the presence of SAM is at least five-fold decreased compared to TrmB D144A, and the apparent association rate is reduced from approximately 60 s^−1^ to 2 s^−1^. As tRNA binding is severely compromised for the TrmB R26A and R155A variants, additional steps in tRNA association are observed in the rapid-kinetic stopped-flow experiments which may reflect conformational events that are typically hidden. In summary, residues R26, T217 and R155 are critically involved in tRNA binding to TrmB. Interestingly, these residues map to three different areas on the surface of TrmB (Figure 1B). As TrmB is a relatively small enzyme of only 27 kDa, our results suggest that TrmB utilizes its extended surface to interact with tRNA. Further structural studies will be needed to uncover the precise orientation of tRNA relative to *E. coli* TrmB.

### Conclusion

Taken together, our results show that TrmB is a biologically-relevant enzyme that confers a fitness advantage to *E. coli* under oxidative stress. As previous reports suggest that this effect may be tRNA specific, it is crucial to understand the interaction of TrmB with tRNA. As a first critical step in this direction, we established a new rapid-kinetic assay complementing standard equilibrium binding assays such as nitrocellulose filtration. Thereby, we have dissected the molecular mechanism of TrmB. Our data suggest that binding of SAM to TrmB induces a conformational change that is required for fast and tight tRNA binding. Three residues located at different areas on the TrmB surface (R26, T217, and R155) are critically contributing to the stable and fast binding of tRNA to this enzyme. Following tRNA binding, the formation of m^7^G46 is presumably the rate-limiting step which is followed by rapid release of the modified tRNA. This insight into the molecular mechanism of TrmB thus lays the ground for future studies to address the role of TrmB-mediated modification of different tRNAs and for structural studies of the TrmB-tRNA complex.

## Experimental procedures

### Buffers and reagents

Experiments were performed in TAKEM_4_ buffer (50 mM Tris-HCl pH 7.5, 70 mM NH_4_Cl, 30 mM KCl, 1 mM EDTA, 4 mM MgCl_2_). [C5-^3^H]UTP for in vitro transcription of [^3^H]tRNA^Phe^ was purchased from Moravek Biochemicals. 5-(iodoacetamido)fluorescein (5-IAF) was from Sigma-Aldrich. SAM was purchased from New England Biolabs. All other chemicals were purchased from Thermo Fisher Scientific.

### Hydrogen peroxide growth comparison for ΔtrmB and its parental strain

The identity of the *ΔtrmB* strain from the Keio collection was validated by colony PCRs targeting the region upstream (5′-GCTGCAACTTCCTCAAAGG-3′) and downstream (5′-CGTCACTGAAAGTGCTGCC-3′) of the *trmB* locus and the kanamycin cassette (k1/k2) (46). For growth analysis, four biological replicates for each the *ΔtrmB* strain and its parental strain, BW25113, were grown overnight in 3-5 mL LB in the presence of 50 µg/mL kanamycin (*ΔtrmB*) or in the absence of antibiotic (wildtype). Cells were resuspended in fresh LB medium lacking antibiotic and diluted to an OD_600_ of 0.1 in 150 µL LB containing 5 mM H_2_O_2_. Cultures were incubated at 37°C for 48 hours with continuous shaking, and the absorbance was measured every 15 minutes at 600 nm in an Eon BioTek 96-well plate reader. The average OD_600_ reading of at least three biological replicates for each strain was plotted versus time with error bars representing the standard error of the mean (SEM).

### Site directed mutagenesis to prepare TrmB variants

Expression plasmids for TrmB variant expression were prepared from the pET28a-TrmB plasmid (29) using Quikchange site-directed mutagenesis with the following overlapping primers:

TrmB D144A: 5′-TTTTCCCTGCCCCGTGGCACAAAGCGC-3′ and 5′-GCCACGGGGCAGGGAAAAAGAGCTGC-3′

TrmB T217A: 5′- CCGGTGGCGAAATTTGAACAACGTGG-3′ and 5′- CAAATTTCGCCACCGGACGTGATGCC-3′

TrmB R26A: 5′- TTTGTGCGCGCCCAGGGGCGACTGAC-3′ and 5′- CCCCTGGCGGCGCACAAAACTACGG-3′

TrmB R155A: 5′- AATAAACGCGCTATCGTTCAGGTGCCG-3′ and 5′- GAACGATAGCGCGTTTATTATGGCGCGC-3′

The sequences of pET28a-TrmB D144A, pET28a-TrmB R26A, pET28a-TrmB R155A, pET28a-TrmB T217A were confirmed by Sanger sequencing (Genewiz).

### Protein expression and purification

Plasmids encoding wildtype or variant TrmB were transformed into BL21 (DE3) cells. Similarly, plasmids pBH113 and pBH402 for ThiI and IscS overexpression were transformed into BL21 (DE3) for overexpression as described in (29). In brief, for overexpression, cells were grown at 37°C in LB broth supplemented with 50 µg/mL kanamycin or 100 µg/mL ampicillin. Protein overexpression was induced at an OD_600_ of approximately 0.6 with 1 mM isopropyl β-d-1-thiogalactopyranoside (IPTG). After three hours, cells were collected by centrifugation at 5000*g* for fifteen minutes, flash frozen, and stored at -80°C.

Proteins were purified via their amino-terminal hexahistidine tag using nickel-sepharose followed by Superdex 75 (XK 26/100) chromatography as previously described (45). TrmB protein concentrations were determined by absorbance at 280 nm using an extinction coefficient of 27,960 M^−1^ cm^−1^ (calculated using ProtParam (47)). Similarly, ThiI and IscS concentrations were determined by A_280_ using experimentally determined extinction coefficients 63,100 and 25,400 M^−1^ cm^−1^, respectively (48). Protein concentrations were validated using comparative SDS-PAGE and Bradford assays.

### tRNA^Phe^ preparation

*E. coli* tRNA^Phe^ was prepared as described previously (45) by amplification from the pCF0 plasmid (49) followed by *in vitro* transcription. The resulting tRNA^Phe^ was purified by phenol extraction and Superdex 75 (XK 26/100) chromatography. For uniformly tritium-labeled tRNA^Phe^, [5-^3^H]UTP was included in the reaction and [5-^3^H]tRNA^Phe^ was purified by Nucleobond Xtra Midi anion exchange gravity columns (Machery-Nagel) (45).

### tRNA^Phe^ modification and fluorescent labeling

In order to introduce a specific fluorescent label at position 8 of the tRNA structure, purified ThiI and IscS enzymes were used to prepare s^4^U8-tRNA^Phe^. For this reaction, tRNA^Phe^ was first folded in TAKEM_4_ buffer by heating to 65°C for five minutes and cooling to room temperature for at least ten minutes. Folded tRNA^Phe^ at a final concentration of 5 µM was incubated with 3 µM IscS and 3 µM ThiI in the presence of 40 µM pyridoxal-5′-phosphate (PLP), 4 mM adenosine triphosphate (ATP), 500 µM L-cysteine, 0.07 U/µL RiboLock RNase inhibitor (ThermoScientific), and 1 mM dithiothreitol in TAKEM_4_ buffer for at least two hours at 37°C in a 20 mL total volume. The reaction was stopped by enzyme denaturation via heating at 80°C for fifteen minutes and enzymes were then removed by phenol/chloroform extraction. Following isopropanol precipitation to reduce the volume, reaction cofactors were removed by Superdex 75 (10/300 GL) chromatography. tRNA thiolation was validated to be at least 80% by ImageJ analysis of 5-100 pmol of tRNA on a 20 µM [(N-acryloylamino)phenyl]mercuric chloride (APM), 7 M urea, 10% polyacrylamide gel (50).

Fluorescent labeling of the s^4^U8 residue was achieved by incubating 60 µM s^4^U8-tRNA^Phe^ with 3.2 mM 5-(iodoacetamido)fluorescein (5-IAF) in 12 mM HEPES-KOH pH 8.2 containing 80% (v/v) dimethyl sulfoxide (DMSO) at 65°C in the dark for four hours, similar to (34). To remove unincorporated dye, at least eight successive phenol extractions were performed until the organic layer was no longer yellow. To remove phenol, two chloroform extractions were performed followed by ethanol precipitation. The final fluorescein-s^4^U8-tRNA^Phe^ was resuspended in water, and the tRNA concentration was determined spectrophotometrically at 260 nm. The labeling efficiency was determined by absorbance at 492 nm, and was typically ~2-15%.

### Stopped-flow to monitor TrmB–tRNA^Phe^ binding

Following tRNA folding, fluorescein-s^4^U8-tRNA^Phe^ (final concentration: 1 µM) was rapidly mixed with TrmB at a final concentration between 3 – 20 µM in TAKEM_4_ buffer in a KinTek SF-2004 stopped flow apparatus at 20°C. Relative fluorescence (*Y*) was plotted against time (*t*), and traces were fit to a one-, two-, or three-exponential function to determine apparent rates (*k*_app_) using TableCurve 2D:

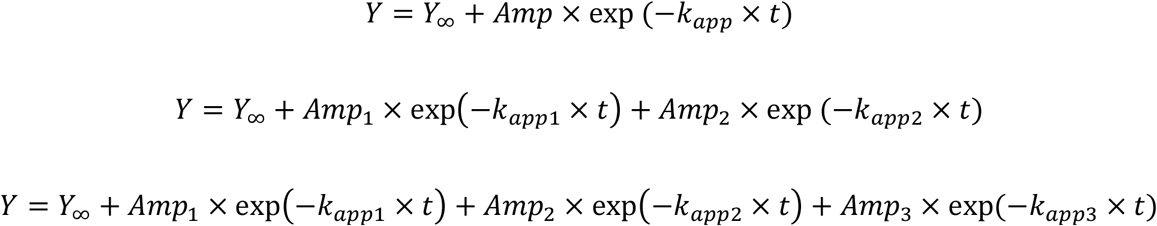

Data shown are averages of at least eight independent time courses.

For TrmB D144A, apparent rates were plotted against enzyme concentration and fit with a linear equation to determine the association rate constant *k*_1_:

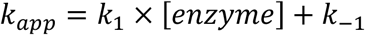

### Nitrocellulose filtration to determine affinity of TrmB for tRNA

Prior to the reaction, [^3^H]tRNA^Phe^ was refolded in TAKEM_4_ buffer by heating and cooling as described above. Refolded [^3^H]tRNA^Phe^ (40 nM) was incubated with increasing concentrations of TrmB wildtype or variant enzyme (0 – 30 µM) in TAKEM_4_ buffer in the presence or absence of 50 µM SAM for ten minutes at room temperature.

As described in (45), the enzyme–tRNA mixture was filtered through a nitrocellulose membrane under vacuum and the proportion of bound tRNA was determined by scintillation counting. To determine the dissociation constant (*K*_*D*_), percent tRNA bound (*Bound*) was plotted as a function of enzyme concentration ([*enzyme*]) and fit with a hyperbolic equation using GraphPad Prism software:

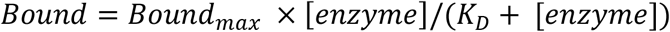

## Data availability

All data are contained within the manuscript.

## Acknowledgements

We thank Saskia Funk for initial fluorescent tRNA labeling and initial stopped-flow experiments, Dr. Eugene Mueller for providing ThiI and IscS expression plasmids, and the National BioResource Project (National Institute of Genetics, Japan) for the *ΔtrmB E. coli* strain from the Keio collection.

## Funding and additional information

This work was supported by the Natural Sciences and Engineering Research Council of Canada [UK: Discovery Grant RGPIN-2020-04965 and Discovery Accelerator Supplement RGPAS-2020-00010].

